# Pneumolysin promotes host cell necroptosis and bacterial competence during pneumococcal meningitis as shown by whole animal dual RNA-seq

**DOI:** 10.1101/2022.02.10.479878

**Authors:** Kin Ki Jim, Rieza Aprianto, Arnau Domenech, Jun Kurushima, Diederik van de Beek, Christina M.J.E. Vandenbroucke-Grauls, Wilbert Bitter, Jan-Willem Veening

## Abstract

Pneumolysin is a major virulence factor of *Streptococcus pneumoniae* that plays a key role in interaction with the host during invasive disease. How pneumolysin influences these dynamics between host and pathogen interaction during early phase of central nervous system infection in pneumococcal meningitis remains unclear. Using a whole animal *in vivo* dual RNA-seq approach, we identified pneumolysin-specific transcriptional responses in both *S. pneumoniae* and zebrafish (*Danio rerio*) during early pneumococcal meningitis. By functional enrichment analysis we identified host pathways known to be activated by pneumolysin, and discovered the importance of necroptosis for host survival. Inhibition of this pathway using the drugs necrostatin-5 or GSK’872 increased host mortality during pneumococcal meningitis. On the pathogen’s side, we find that pneumolysin-dependent competence activation is crucial for intra-host replication and virulence and that not all bacteria activate competence at the same time. Altogether, this study provides new insights into pneumolysin-specific transcriptional responses and identifies key pathways involved in pneumococcal meningitis.

**GRAPHICAL ABSTRACT:** 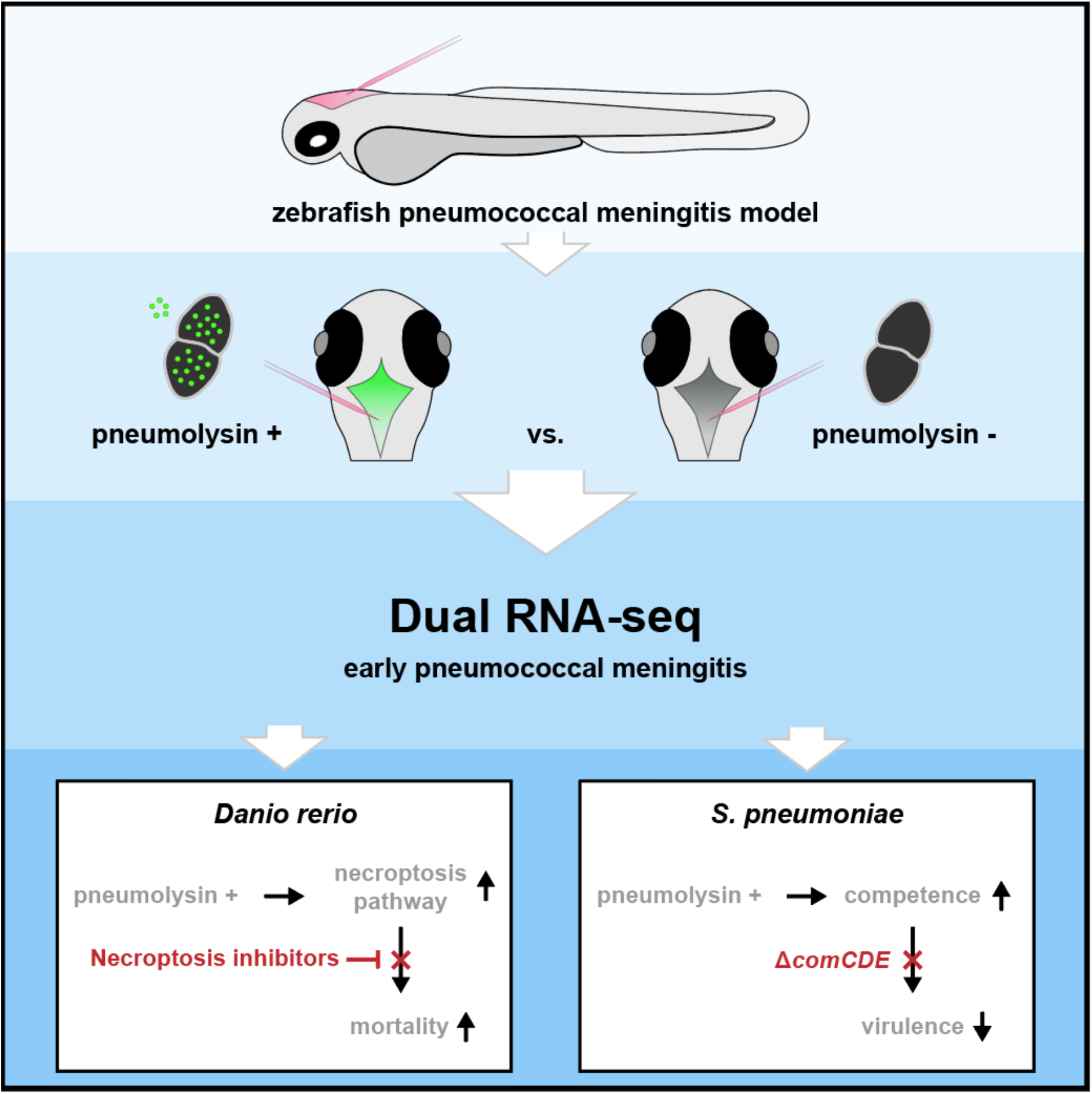

**HIGHLIGHTS:**

- Pneumolysin-specific host and bacterial responses as identified by whole animal dual RNA-seq, available at https://veeninglab.com/dual-danio
- Discovery of a functional necroptosis or necroptosis-like pathway in zebrafish
- Heterogeneity in competence development during infection
- Competence development is an important virulence determinant

## INTRODUCTION

The interaction between host and microbe determines the pathogenesis of infectious diseases. In invasive pneumococcal disease (IPD), caused by the opportunistic human pathogen *Streptococcus pneumoniae*, pneumococci need to successfully colonize mucosal surfaces of the nasopharynx by adapting to the host environment, before they can spread into the blood stream, lungs or central nervous system and cause IPD. Upon invasion, the pathogen will be detected by pattern recognition receptors (PRRs), leading to the activation of the innate immune response that will try to eliminate the pneumococci. In turn, pneumococci will counteract by employing a variety of strategies to escape from eradication by the host’s immune response (van de Beek et al., 2021; Henriques-Normark and Tuomanen, 2013; Kadioglu et al., 2008; Weiser et al., 2018). Insight into these complex host-pneumococcus interaction dynamics may lead to better understanding in the factors leading to poor disease outcome and to develop new intervention strategies (van de Beek, 2012). Despite the introduction of pneumococcal conjugate vaccines, the global burden of pneumococcal disease remains high, with an estimated 3.7 million episodes and up to half a million deaths every year in children under 5 years alone (Wahl et al., 2018).

To study infection dynamics, simultaneous profiling of both host and pathogen transcriptional responses, so-called dual RNA-seq, has been adopted as a tool which offering limited technical bias and higher efficiency as compared to conventional approaches (single species approach, array-based methods) (Westermann and Vogel, 2018; Westermann et al., 2016, 2017; Wolf et al., 2018). In previous studies, using dual RNA-seq, we showed that *S. pneumoniae* represses the innate immune response in human alveolar lung epithelial cells and activates its own mucin-dependent sugar transporters upon adherence (Aprianto et al., 2016). This was also shown in a murine pneumonia model (Minhas et al., 2020). So far, dual RNA-seq has mainly been applied in *in vitro* models of bacterial infection, which has led to valuable new insights into host-pathogen interactions for pathogens such as *Staphylococcus aureus, Plasmodium* spp., *Pseudomonas aeruginosa, Salmonella* spp., and *S. pneumoniae* (Aprianto et al., 2016; Baddal et al., 2015; Dillon et al., 2015; Nuss et al., 2017; Rosenberg et al., 2021; Thänert et al., 2017; Tierney et al., 2012; Westermann et al., 2016, 2019). More recently, this approach has been also employed in *in vivo* infection models (Damron et al., 2016; Kumar et al., 2018; LaMonte et al., 2019; Zhang et al., 2019), although most of these experiments were often combined with some (flow-) sorting steps to identify infected cells. Recently, the *in vivo* pneumococcal and mouse transcriptomes within the nasopharynx, lungs, blood, heart, and kidneys were mapped revealing distinct gene-expression profiles depending on organ and disease state (D’Mello et al., 2021). Together, these studies have demonstrated that dual RNA-seq can provide invaluable insights into host-microbe interactions.

Here we performed, to the best of our knowledge, the first whole animal *in vivo* dual RNA-seq study. We applied whole animal *in vivo* dual RNA-seq to specifically examine the role of pneumolysin on the transcriptional response of both pathogen (*S. pneumoniae* D39V) and host (*Danio rerio*) during early pneumococcal meningitis in a zebrafish infection model (Jim et al., 2016). Pneumolysin, a major pneumococcal virulence factor, is a cholesterol-dependent pore-forming cytolysin produced by all known clinical isolates (Kadioglu et al., 2008; Weiser et al., 2018). It has been shown to play a role in the pathogenesis and pathophysiology of pneumococcal meningitis (Mook-Kanamori et al., 2011; Surve et al., 2018) and infection with pneumolysin-deficient mutants causes mild to no disease in experimental pneumococcal meningitis models (Hirst et al., 2008; Jim et al., 2016; Wellmer et al., 2002). Pneumolysin might also play a key role in transmission by stimulating the host immune response (Weiser et al., 2018). Moreover, persisting high levels of pneumolysin in the cerebrospinal fluid of patients with pneumococcal meningitis is associated with mortality (Wall et al., 2012). Pneumolysin was not required for pneumococcal replication in a murine influenza A virus-superinfection model of pneumonia (Liu et al., 2021), further highlighting our knowledge gap related to pneumolysin and its role in pathogenesis.

The serotype 2 *S. pneumoniae* strain D39 is one of the most used strains to study pneumococcal biology and infection. The genome of this strain, and a well-defined genome-wide transcriptional response, have been annotated in detail (Aprianto et al., 2018; Slager et al., 2018). On the host side, the zebrafish has emerged as an *in vivo* model for various infections, including pneumococcal infections (Jim et al., 2016; Lieschke and Currie, 2007; van der Sar et al., 2004). Zebrafish have several advantages over mice, including small size and high fecundity; zebrafish embryos develop extra maternally and are transparent, which makes them highly suitable for whole-organism based high-throughput screening (Lieschke and Currie, 2007; van der Sar et al., 2004). At a low dose of infection (300 CFU of pneumococci), zebrafish embryos develop disease progression and ultimately succumb to meningitis (Jim et al., 2016). In the present study, we identified known pathways activated by pneumolysin and novel pneumolysin-specific host-pathogen interactions in early pneumococcal meningitis, thereby providing a valuable resource for future studies. The data is easily accessible, searchable, and downloadable via a web-based platform (https://veeninglab.com/dual-danio). In this study, we demonstrated that pneumococcal competence-activation is a key hallmark of disease progression and in meningitis delineate the essential role of pneumolysin in this process.

## RESULTS

### Dual RNA-seq in early pneumococcal meningitis in zebrafish larvae

To study host-pathogen interaction in early pneumococcal meningitis, we used our zebrafish embryo meningitis model and dual-RNA seq pipeline, and performed RNA-seq simultaneously on both *D. rerio* and *S. pneumoniae* (Aprianto et al., 2016; Jim et al., 2016). To determine pneumolysin-specific transcriptional changes we injected 2 days post-fertilization (dpf) zebrafish larvae with a pneumolysin-deficient *S. pneumoniae* mutant strain (PLY-) and its complemented version (PLY+), in which the deleted pneumolysin gene is restored and thus able to produce pneumolysin (Figure 1A). Survival experiments showed that *S. pneumoniae* PLY-was significantly attenuated as compared to *S. pneumoniae* PLY+ in the zebrafish meningitis model, similar to what we have previously reported (Jim et al., 2016; Figure S1). Next, we wanted to determine the specific contributions of pneumolysin to transcription changes in the host and the pathogen. Considering the size of the host and the relatively small pathogen burden, the RNA signal will be dominated by *D. rerio* RNA. Therefore, we increased the inoculum of *S. pneumoniae* threefold (to 2000 CFU) as compared to our previous work. Three biological replicates per group were used, with 100 larvae pooled per replicate (total n = 300 per group). Importantly, at 8 hours post injection (hpi), the estimated total amount of pneumococci per group per replicate was comparable between both groups (∼2 million pneumococci per replicate; *p* value = 0.3281) (Figure 1B). To minimize transcriptional changes caused by sample handling, we did not separate pneumococci from *D. rerio* cells before total RNA isolation and dual rRNA depletion (Figure 1C). The obtained dual RNA-seq dataset was pruned and handled as described previously (Figure 1D) (Aprianto et al., 2016). In short, raw reads were trimmed and aligned to a chimeric concatenated genome of *D. rerio* and *S. pneumoniae*. After alignment, one-step mapping was performed to minimize false negatives. The resulting raw counts were counted separately and categorized as either *D. rerio* (ENSEMBL, release 11) or pneumococcal (D39V, Slager et al., 2018). Differential gene analysis was calculated by DESeq2 and specific groups of genes were removed from subsequent analysis (Love et al., 2014). Finally, unbiased automated clustering (Kumar and Futschik, 2007), and functional enrichment analysis were performed (Carbon et al., 2009; Liao et al., 2019; Subramanian et al., 2005).

**Figure 1.**
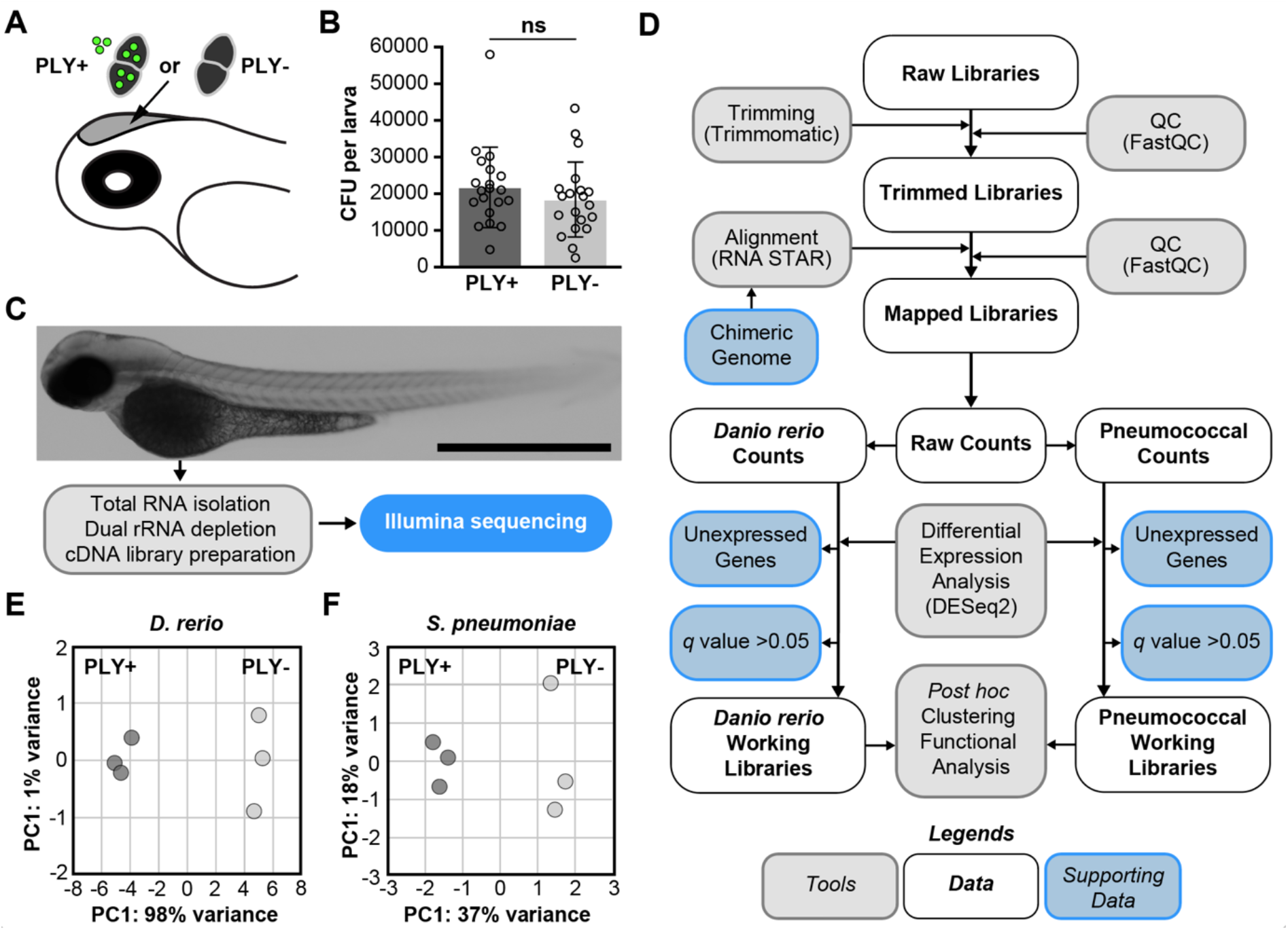
Dual-RNA sequencing of host-pathogen transcriptomes in early pneumococcal meningitis. **(A)** Zebrafish meningitis infection model. Pneumolysin-deficient *S. pneumoniae* D39V (PLY-) mutant bacteria or complemented *S. pneumoniae* D39V (PLY+) bacteria (∼2.000 CFU) were injected into the hindbrain ventricle of 2 days post fertilization zebrafish larvae. **(B)** Bacterial load in zebrafish larvae infected with *S. pneumoniae* PLY+ or *S. pneumoniae* PLY-at 24 hours post injection. The data represents the mean ± SD of two individual experiments; each dot represents a single larva; ns: non-significance; determined by unpaired t-test. **(C)** Total RNA was isolated from pooled infected zebrafish larvae (n=100 per replicate) for preparation of cDNA libraries and sequencing at 8 hours post injection. **(D)** Quality Control (QC) was performed on the raw reads, low-quality reads trimmed, and the remaining reads aligned to a synthetic chimeric genome. Aligned reads were counted and classified as pneumococcal or *Danio rerio* counts. Final working libraries were created after removal of two gene fractions, and clustering and functional enrichment analysis were performed. **(E)** Principal Component (PC) Analysis plot of pathogen transcriptional response to infection showed that the replicates cluster closely together. **(F)** PC Analysis plot of host response showed similar clustering behavior.

Global descriptive analysis showed that the sizes of the six dual RNA-seq libraries were well-balanced (average 70 million reads, range: 55 to 99 million reads, Figure S3A). Moreover, 99.94% of reads could be aligned onto the zebrafish genome (99.94 – 99.95%) with the remaining of the reads mapping to the pneumococcal genome. The number of reads translate into an average depth of 4.2 (3.3 to 5.9 times) for the host and 1.53 for the pneumococcal libraries (1.22 to 2.35 times) (Figure S3B). In addition, for all datasets, principal component (PC) analyses showed that the replicates cluster close together (Figure 1E-F), indicating the homogeneity of response within the replicates and the transcriptional dissimilarity because of the presence or absence of pneumolysin. In the zebrafish dataset, there were 2,230 genes out of 25,592 (69%) in the raw library mapped (Figure 1G, Table S1). After removal of unexpressed genes and genes with no significant difference (*p* value < 0.05, *q* value by DESeq2), the zebrafish working libraries contained 6,403 genes (25%). In the pneumococcal libraries, out of 2,133 annotated genes, 1,924 genes (89%) were filtered in and fed into downstream analysis (Figure 1G, Table S2,). The excluded 201 genes were not further processed because their mean normalized counts were less than the optimal threshold, as described by DESeq2 (Love et al., 2014).

The raw data is publicly available at the Gene Expression Omnibus (GEO) under accession number GSE123988. To make the data more easily accessible, we also host the complete dual RNA-seq database online (https://veeninglab.com/dual-danio). On this website users can select the gene of interest and download their expression level (Figure S4).

### Pneumolysin-specific host responses in pneumococcal meningitis

In the host transcriptional response, approximately 25% of the total zebrafish genes were differentially expressed at 8 hours post injection of *S. pneumoniae* PLY+ as compared to zebrafish infected with *S. pneumoniae* PLY-. More specifically, 3,753 (14.7 %) transcripts were significantly more abundant in zebrafish larvae infected with *S. pneumoniae* PLY+, whereas 2,650 genes (10.4%) were more abundant in zebrafish infected with *S. pneumoniae* PLY- (*q* value < 0.05) (Figure 2A). Gene expression values of five selected host transcripts were confirmed by RT-qPCR (Figure 2B). After applying a fold change (FC) cut-off at 1.5 for all comparisons, we identified 341 (1.3%) genes that are differentially expressed between both groups. The application of this more stringent cut-off resulted in a major shift of differentially expressed genes towards *S. pneumoniae* PLY+ infected larvae; in total 312 genes (91.5%) were enriched in larvae infected with *S. pneumoniae* PLY+ as compared to 29 (8.5%) genes in larvae infected with *S. pneumoniae* PLY- (Figure 2A and Table S1).

**Figure 2.**
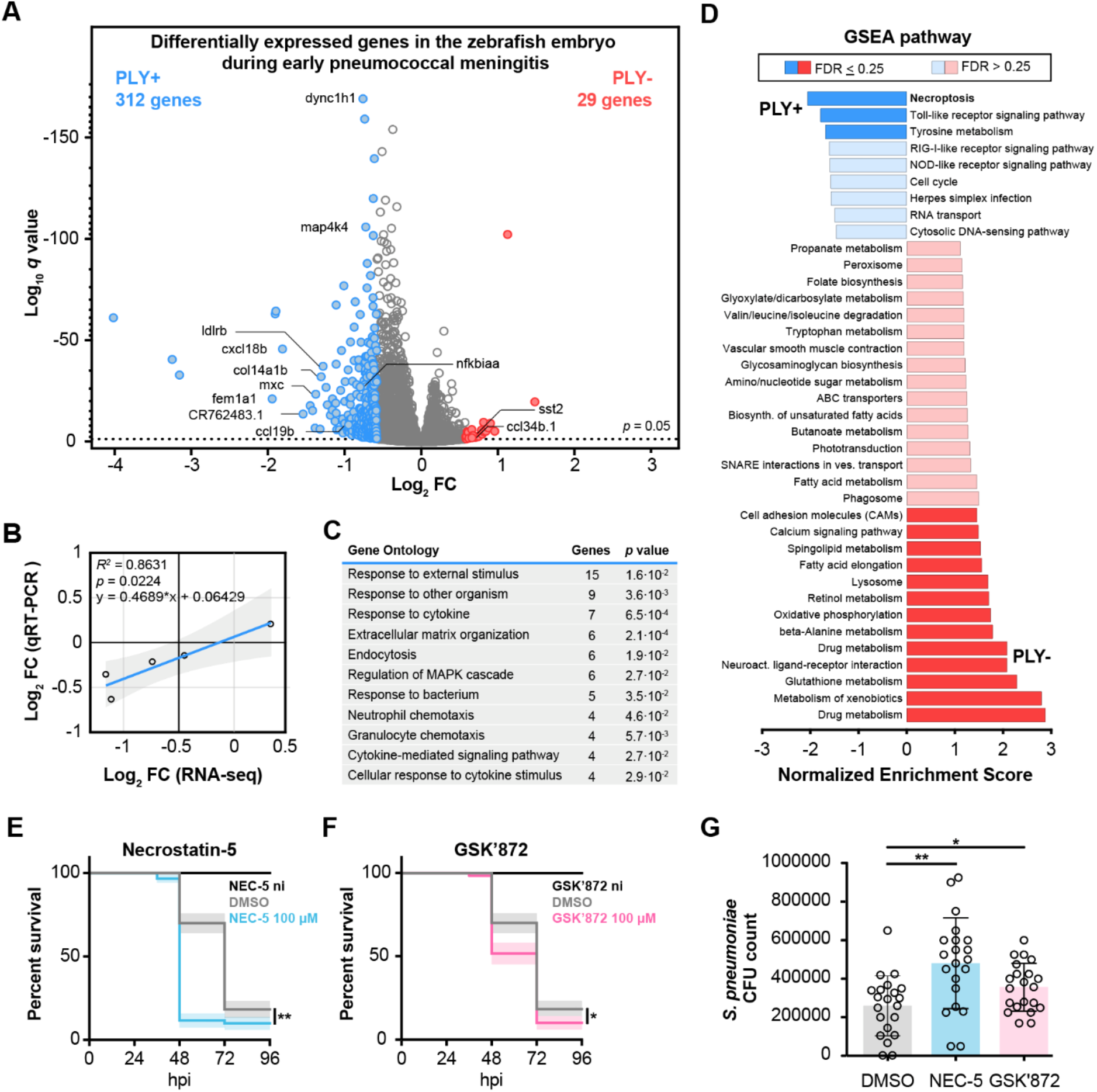
The host’s transcriptional response to infection with pneumolysin-positive or pneumolysin-negative *S. pneumoniae* D39V. **(A)** Volcano plot from DESeq2 analysis of RNA pools from zebrafish larvae infected with the pneumolysin-deficient *S. pneumoniae* D39V mutant strain (PLY-) or the complemented *S. pneumoniae* D39V mutant strain (PLY+) at 8 hours post infection. The presence of pneumococcal pneumolysin activates a multitude of host genes in response to *S. pneumoniae* infection (FC > 1.5, *q* value < 0.05). (**B**) Gene expression correlation between RNA-seq and RT-qPCR data; each dot represents a single gene; correlation co-efficient (Pearson) and linear regression are indicated. **(C)** Gene Ontology term enrichment analysis after DESeq2 analysis showed significant enrichment of several immunomodulatory pathways. Note that an individual gene can be part of multiple terms. (**D**) Gene Set Enrichment analysis after DESeq2 analysis showed enrichment of several immunomodulatory pathways in the presence of pneumolysin, whereas the phagolysosome pathway and many metabolic pathways were enriched in the absence of pneumolysin (FDR <0.50). **(E, F)** Survival curves of 2 dpf zebrafish larvae injected with *S. pneumoniae* D39V in the hindbrain ventricle and control non-infected (ni) zebrafish larvae treated with necrostatin-5 (RIPK1 inhibitor), GSK’872 (RIPK3 inhibitor) or vehicle (DMSO). Larvae were infected with 400 CFU. The data represent the mean ± SEM of three individual experiments with 20 embryos in each group; **(E)** *p* value < 0.0001, vehicle-treated versus necrostatin-5 (100 μM), **(F)** *p* value < 0.0001, vehicle-treated versus GSK’872 (100 μM); determined by log-rank test. **(G)** Bacterial loads at 24 hours post injection with *S. pneumoniae* D39V. Treatment with necrostatin-5 or GSK’872 resulted in a higher bacterial load. Larvae were infected with 1.000 CFU. The data represent the mean ± SEM of two individual experiments; *p* value = 0.0011, vehicle-treated (DMSO) versus necrostatin-5 (100 μM); *p* value = 0.004, vehicle-treated (DMSO) versus GSK’872 (100 μM); determined by Mann-Whitney U test.

### Functional enrichment analyses identify several host pathways involved in pathogen detection and clearance

To analyze the differentially expressed genes, we performed functional enrichment analyses to determine their significance in the context of the background set of genes. First, we performed Gene Ontology (GO) biological process enrichment analysis (cut-off > 3 genes, *p* value < 0.05) on significantly enriched genes (*q* value < 0.05, FC > 1.5) in the presence (n = 312) or absence (n = 29) of pneumolysin to identify which GO terms were over-represented. In the presence of pneumolysin, GO enrichment analysis mainly identified genes known to be involved in immune signaling, including “response to cytokine” (*p* value = 6.5 × 10^−3^), “neutrophil chemotaxis” (*p* value = 4.6 × 10^−2^), “granulocyte chemotaxis” (*p* value = 5.7 × 10^−3^), “cytokine-mediated signaling pathways” (*p* value = 2.7 × 10^−2^) and “cellular response to cytokine stimulus” (*p* value = 2.9 × 10^−2^) (Figure 2C, Table S3). These genes encode for chemokines (Ccl19b, Cxcl18b), cytokine controlling transcription factors (June, Junba, Junbb), interleukin receptor (Il10rb) and other immunomodulatory proteins (Irak3, Mmp9, S1pr4, Tab1) (Table S1). Other enriched GO terms include “extracellular matrix structure” and “extracellular matrix organization”, which have been suggested to play a role in signaling and coordination of the migration of leukocytes, thus contributing to the inflammatory response against infections (Tomlin and Piccinini, 2018; Vaday and Lider, 2000). In the absence of pneumolysin we identified genes involved in terms associated with catabolic processes (Table S3).

Next, we performed Gene Set Enrichment Analysis (GSEA) at a low statistical cutoff (FDR < 0.50) to evaluate the differential gene expression data at the level of gene sets and to gain a rough insight in biologically relevant pathways and processes (Subramanian et al., 2005). GSEA also analyses differentially expressed and functionally annotated genes in a dataset, but discards the fold-change cut-off threshold limitation and uses a ranked list based on the direction and magnitude of expression change of these genes (Subramanian et al., 2005). Pathways known to be involved in intrinsic immune host defense mechanisms against pathogens were mainly enriched in zebrafish embryos infected with PLY+, including pattern recognition receptor (PRR) pathways, such as the toll-like receptor (TLR) signaling pathway (FDR = 0.087), RIG-I-like receptor (RLR) signaling pathway (FDR = 0.343), and NOD-like receptor signaling pathway (FDR = 0.305) (Figure 2D, Table S3). At this stage of infection, the number of bacteria is comparable between both groups, suggesting that the enrichment of these pathways is due to the presence of pneumolysin (Figure 1B). PRRs are important receptors that recognize both exogenous as well as endogenous ligands, thereby contributing to the recognition of molecular structures of invading pathogens by the innate immune system (Takeuchi and Akira, 2010). In the absence of pneumolysin, we identified many metabolic pathways, including fatty acid metabolism (FDR = 0.260), oxidative phosphorylation (FDR = 0.123), and glutathione metabolism (FDR = 0.004). Moreover, phagosome (FDR = 0.260) and lysosome (FDR = 0.131) pathways were enriched in larvae infected with PLY-, suggesting more active phagocytosis by the host in the absence of pneumolysin (Figure 2D, Table S3).

### Necroptosis pathway contributes to pathogen clearance

A highly enriched immunoregulatory pathway in the presence of pneumolysin, that was less expressed in larvae challenged with the pneumolysin mutant, was the necroptosis pathway (FDR = 0.002) (Figure 2D, Table S3). Necroptosis is a caspase-independent form of programmed cell death that depends on activation of the key molecules receptor-interacting protein kinase 1 (RIPK1), receptor-interacting protein kinase 3 (RIPK3) and mixed lineage kinase domain-like (MLKL), and has been shown to regulate inflammation (Kearney and Martin, 2017; Linkermann and Green, 2014; Vandenabeele et al., 2010). Both *ripk1l* (FC −1.09, *q* value = 0.364) and *ripk3* (FC −1.08, *q* value = 0.807) were enriched in the presence of pneumolysin-expressing bacteria, albeit not statistically significantly. For MLKL, we could not identify a zebrafish ortholog (see Discussion). To study the role of this necroptosis or necroptosis-like pathway in pneumococcal meningitis, we first infected zebrafish larvae with wild type *S. pneumoniae* D39V in the hindbrain ventricle and treated them with necroptosis pathway inhibitors necrostatin-5, a RIPK1 inhibitor, or GSK’872, a RIPK3 inhibitor (Degterev et al., 2008; Mandal et al., 2014). As shown in Figure 2E-G, inhibition of the necroptosis pathway by treatment with 100 μM necrostatin-5 or 100 μM GSK’872 resulted in a higher mortality rate and higher bacterial load in zebrafish larvae as compared to DMSO treatment. Treatment of non-infected zebrafish with necrostatin-5 or GSK’872 alone did not result in a higher mortality rate, suggesting that the inhibitor itself had no toxic effect at this concentration (Figure 2E-2F). Together, these data suggest that necroptosis plays a protective role in pneumococcal meningitis in our model.

### Pneumolysin contributes to chemokine and cytokine signaling during infection

Among the most highly enriched transcripts in the presence of pneumolysin we identified genes directly or indirectly involved in chemokine and cytokine signaling (e.g. *fem1a, plxnd1*, ENSDARG00000114454 (growth-regulated alpha protein-like), *ldlrb, mxc*, and *cxcl18b*), confirming the role of pneumolysin as an important immunomodulatory virulence factor (Figure 2A, 3A) (Kadioglu et al., 2008; Kroetz et al., 2015; Netea et al., 1996; Shimizu et al., 2013; Takayama et al., 2006; Torraca et al., 2017). In the presence of pneumolysin, the most strongly enriched gene with known function was *fem1a* (FC −3.85, *q* value = 7.55 × 10^−22^) (Figure 2A, 3A), also named EP4 receptor-associated protein. This protein is known to interact with nuclear factor NF-κB and suppresses inflammatory activation in human macrophages, but also promotes inflammatory activation of murine microglia cells through mitogen-activated protein kinase 4-mediated signaling (Fujikawa et al., 2016; Minami et al., 2008; Takayama et al., 2006). Strikingly, in our dataset NF-κB variants (*nfkbiaa, nfkbiab, nfkb1, nfkbz*), and *mapk4* were also significantly enriched in the presence of pneumolysin-expressing bacteria (Table S1). Another enriched gene is *cxcl18b* (FC −2.16, *q* value = 5.39 × 10^−39^) (Figure 2A, 3A), a piscine- and amphibian-specific chemokine with similar function as Cxcl8a/Interleukin-8, that exerts a neutrophil chemotactic function. Interestingly, these do not represent phagocytic cells, but rather non-infected cells in tissue close to the site of infection (Torraca et al., 2017). To confirm the higher expression of *cxcl18b* in the presence of pneumolysin-positive bacteria, we used the transgenic zebrafish line *Tg(cxcl18b:eGFP)*, which allows visualization of *cxcl18b* expression (Torraca et al., 2017), and compared zebrafish larvae injected with *S. pneumoniae* PLY+ into the hindbrain ventricle with zebrafish larvae injected with a similar dose of *S. pneumoniae* PLY-. Injection of *S. pneumoniae* PLY-bacteria induced *cxcl18b* expression above control levels (injection with just PBS) (Figure 3B, 3C). However, injection with *S. pneumoniae* PLY+ resulted in significantly higher expression of *cxcl18b* in the brain region over time as compared to injection with *S. pneumoniae* PLY- or injection with PBS (Figure 3B, 3C, Movie S1).

**Figure 3.**
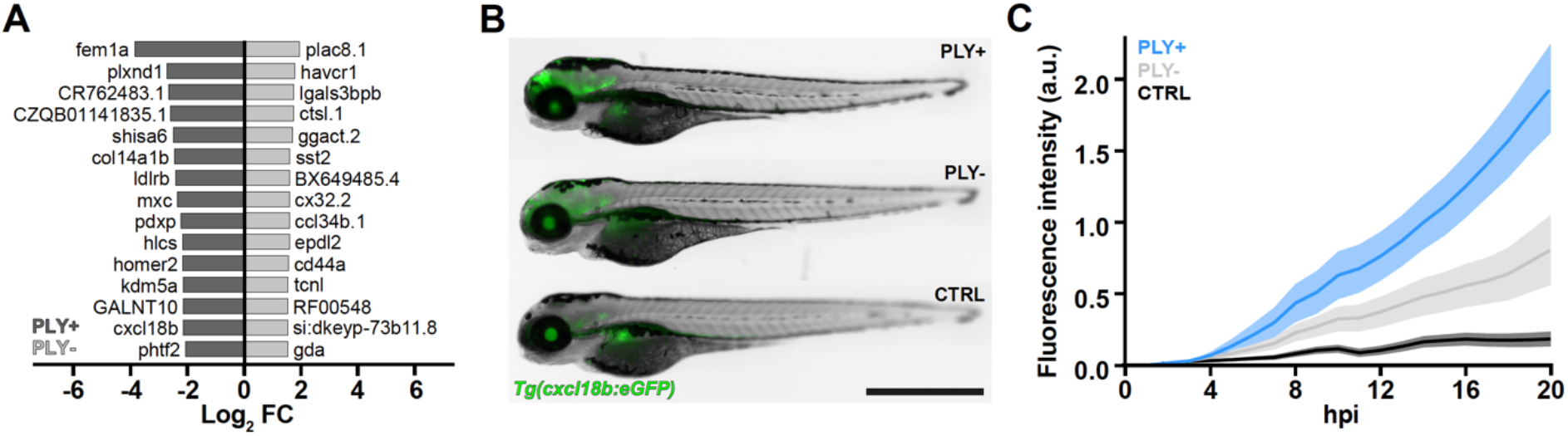
Cxcl18 expression in zebrafish larvae infected with pneumolysin-positive or pneumolysin-negative *S. pneumoniae* D39V. (**A)** Top 20 highest significantly enriched genes in zebrafish infected with *S. pneumoniae* PLY+ or *S. pneumoniae* PLY- (*q* value < 0.05). (**B**) Expression of *Cxcl18b:eGFP* is higher in zebrafish larvae (2 dpf) infected with ∼2,000 colony forming units of *S. pneumoniae* PLY+ as compared to infection with similar numbers of *S. pneumoniae* PLY- or no infection. Scale bar: 500 μm. (**C**) Quantification of *Cxcl18b:eGFP* expression in the head region over time. The data represents the mean ± SEM with 10 embryos in each group; *p* value < 0.0001; determined by one-way ANOVA.

### Pneumolysin-specific pneumococcal transcriptional changes in early pneumococcal meningitis

Due to the relative lower number of reads of the pneumococcal transcriptomes compared to the zebrafish transcriptomes, we applied a softer cut-off (limiting only *p* value, *q* value < 0.5 without fold change limitation) to obtain a more complete impression of the role of pneumolysin during early pneumococcal meningitis (Table S2). Using this cut-off, eight pneumococcal genes were differentially expressed between *S. pneumoniae* PLY+ and *S. pneumoniae* PLY- (*q* value < 0.5); all of them were expressed more in the complemented *S. pneumoniae* PLY+ strain (Figure 4A). The eight genes are coding for: the Clp protease component ClpL; transporters for oligopeptides (AliA) and amino acids (GlnH); enzymes in pyrimidine biosynthesis cascade (PyrB-CarB); competence-induced single-stranded DNA binding protein (SsbB) and two hypothetical proteins (SPV_0145-6), one of which is putatively encoding a metalloproteinase. Since *aliA* is under the regulation of CodY, a global regulator of protein metabolism, and *glnH* is under the regulation of ArgR, an arginine repressor, the complemented *S. pneumoniae* PLY+ strain possibly encounters different nutrients than its cognate pneumolysin deletion *S. pneumoniae* PLY-strain. This hypothesis is further supported by the higher expression of *pyrB-carB*, coding for enzymes in the pyrimidine biosynthesis cascade; both genes are organized in one operon under PyrR regulation, which in turn is sensitive to the levels of intracellular uridine (Figure 4A) (Slager et al., 2018).

**Figure 4.**
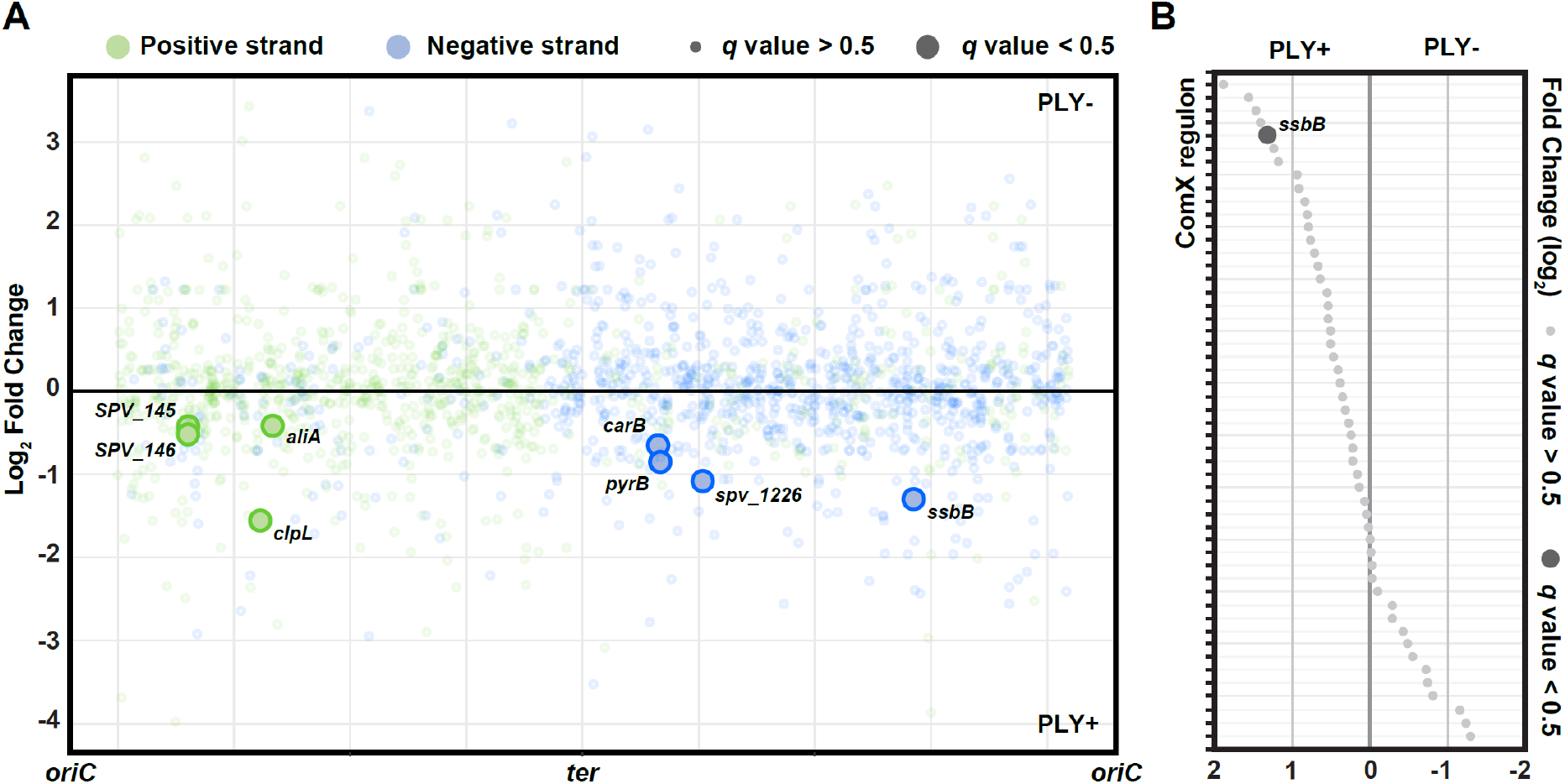
Transcriptional response in pneumolysin-positive or pneumolysin-negative *S. pneumoniae* D39V. **(A)** Pneumolysin-specific transcriptional rewiring in *S. pneumoniae* in response to injection in zebrafish embryo. Fold change of transcriptional response in *S. pneumoniae* PLY+ and PLY-is plotted against its genomic location. (**B)** Fold change of ComX regulons between *S. pneumoniae* PLY+ and *S. pneumoniae* PLY-strains shows gene expression regulation in response to zebrafish infection. *ssbB*, a member of the regulon shows increased expression in *S. pneumoniae* PLY+ as compared to *S. pneumoniae* PLY- (FC > 2, *q* value < 0.5). Regulons are sorted by fold change value in descending order.

The higher expression of *ssbB*, a ComX regulon (Slager et al., 2019), indicates that at least a subset of *S. pneumoniae* PLY+ bacteria is inducing competence genes inside the host. Competence activation is consistent with the expression of *clpL*, as previously observed (Aprianto et al., 2018). In addition to *ssbB*, 37 other competence genes were expressed at higher levels in the complemented *S. pneumoniae* PLY+ strain, albeit not significantly (Fig. 4B). Surprisingly, also a small subset of competence genes (16) was upregulated in pneumolysin-deficient *S. pneumoniae* (Figure 4B).

### Competence development promotes virulence during early meningitis

The *ssbB* gene is under direct control of the competence sigma factor ComX and strongly correlates with the incorporation of external DNA and is a good reporter for competence (Prudhomme et al., 2006; Slager et al., 2014). To monitor whether *ssbB* is activated during early pneumococcal meningitis, we constructed a reporter strain that expresses GFP upon *ssbB* activation (SsbB-GFP). As a control, this strain also constitutively expresses RFP. Next, the competence-reporter strain was injected into the hindbrain ventricle of zebrafish larvae. From as early as ∼2 hpi onwards we observed heterogeneous SsbB-GFP expression, which increases over time in the bacterial population growing at the site of injection, demonstrating that competence is activated in part of the bacterial population during early pneumococcal meningitis (Figure 5A). Interestingly, the expression of SsbB-GFP appears to be induced upon initial contact with phagocytes (Figure 5B). However, at later stages of infection we observed that large numbers of pneumococci were also expressing SsbB-GFP while not being phagocytosed (Figure 5C).

**Figure 5.**
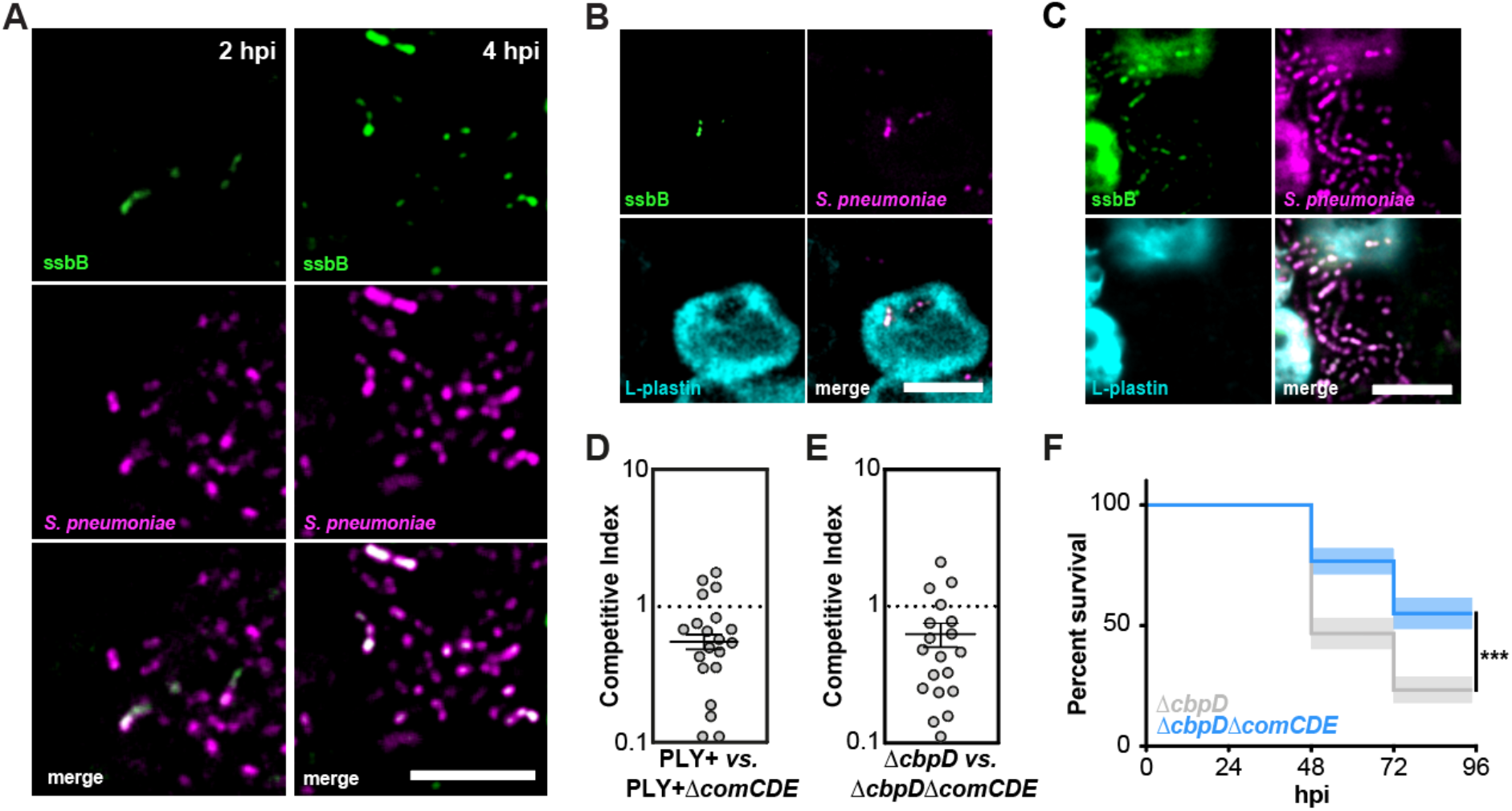
Pneumococcal competence in zebrafish larvae with pneumococcal meningitis. **(A)** Single plane confocal microscopy images showing increased heterogeneous expression of *ssbB*-GFP over time in constitutively HlpA-mKate2 expressing pneumococci injected into the hindbrain ventricle of 2 dpf zebrafish larvae. **(B)** Expression of *ssbB-GFP* upon phagocytosis at 4 hours post injection (hpi) and **(C)** in the absence of phagocytes at 8 hpi. Embryos were injected with ∼2000 CFU. Scale bars: 10 μm. **(D)** Competitive index (CI) analysis of 2 dpf larvae co-injected with similar number of *S. pneumoniae* D39V PLY+ (HlpA-GFP) and *S. pneumoniae* D39V PLY+ Δ*comCDE* (HlpA-mKate2) and **(E)** competitive index of larvae co-injected with similar number of *S. pneumoniae* D39V Δ*cbpD* and *S. pneumoniae* D39V Δ*cbpD*Δ*comCDE*. Larvae were harvested at 24 hpi. A CI score of 1 denotes no difference in virulence. The data represent the mean ± SEM of two individual experiments with 10 larvae in each group; each dot represents a single larva; *p* value <0.0001, D39V PLY+ vs. D39V PLY+ Δ*comCDE; p* value <0.01, D39V Δ*cbpD* vs. Δ*cbpD*Δ*comCDE*; determined by one sample t-test. **(F)** Survival curves of 2 dpf zebrafish larvae injected with 400 CFU of *S. pneumoniae* D39V PLY+ or *S. pneumoniae* D39V PLY+ Δ*comCDE* into the hindbrain ventricle. The data represent three individual experiments with 20 larvae in each group; *p* value = 0.0001; determined by log-rank test.

We hypothesize that competence plays a role in virulence in pneumococcal meningitis. To study whether competence influences virulence in pneumococcal meningitis, we constructed a *comCDE* knock-out mutant strain in the *S. pneumoniae* PLY+ background as this operon controls competence regulation (Johnston et al., 2014). First, we performed a competitive index experiment by co-injecting both wild type (PLY+) and PLY+Δ*comCDE* strains into the zebrafish embryo hindbrain, with a similar number of bacteria and determined the number of bacteria before injection and at 24 hours post injection. We observed that the number of *S. pneumoniae* PLY+Δ*comCDE* (HlpA-mKate2) present in zebrafish larvae at 24 hours post injection was lower as compared to *S. pneumoniae* PLY+ (HlpA-GFP), suggesting that deletion of the *comCDE* operon resulted in decreased fitness as compared to the *S. pneumoniae* PLY+ strain (*p* value<0.0001) (Figure 5D). To exclude the possibility that this decreased fitness is caused by fratricins produced by competent wild type bacteria causing lysis of the Δ*comCDE* mutant (Claverys and Håvarstein, 2007; Shanker and Federle, 2017), we constructed a *S. pneumoniae* D39V Δ*cbpD* mutant that is unable to produce the major fratricin, *i*.*e*., murine hydrolase CbpD. We compared this mutant in a competition experiment with a *S. pneumoniae* D39V Δ*cbpD*Δ*comCDE* double mutant. Consistent with the previous experiment, the number of *S. pneumoniae* D39V Δ*cbpD*Δ*comCDE* was also lower compared to *S. pneumoniae* D39V Δ*cbpD* (*p* value<0.01) (Figure 5E), in line with the observation that competence is induced in spatially clustered foci within the zebrafish (Figure 5A). Additionally, we also performed individual survival experiments with the separate strains. After injection with *S. pneumoniae* D39V Δ*cbpD*Δ*comCDE* we observed a significantly lower larvae mortality rate as compared to injection with *S. pneumoniae* D39V Δ*cbpD* (*p* value = 0.0001) (Figure 5F). Together, these results indicate that the *comCDE* operon and therefore competence is involved in pneumococcal virulence. These results are in line with recent findings in a murine model of pneumococcal meningitis (Schmidt et al., 2019).

## DISCUSSION

We present a detailed dataset of pneumolysin-specific zebrafish-pneumococci transcriptional responses in early pneumococcal meningitis using whole animal *in vivo* dual RNA-seq. This whole animal approach alleviates the need for dissecting and/or sorting of infected tissues thereby reducing experimental handling and technical noise. Our experimental setup allowed us to identify not only previously reported, but also novel pneumolysin-specific transcriptional responses in both host and pathogen. We found that in the presence of pneumolysin, major immunomodulatory pathways were enriched and that competence plays a role in pneumococcal virulence in pneumococcal meningitis among others.

Activation of the immune system in the cerebrospinal fluid is dependent on PRRs (van de Beek et al., 2016; Mook-Kanamori et al., 2011). In our study, we found that the TLR signaling pathway, RIG-I-like receptor pathway, and NOD-like receptor signaling pathway were enriched in zebrafish larvae infected with pneumococci that produce pneumolysin (Figure 2D, Table S2). Malley et al. have shown that in mice the TLR4 signaling pathway is responsible for recognition of pneumococcal pneumolysin (Malley et al., 2003). In zebrafish, however, there is evidence that TLR4 proteins do not function in a similar fashion as in mammals (Sepulcre et al., 2009; Sullivan et al., 2009). The only *tlr4*-like gene we found significantly enriched was *tlr4ba* (FC 1.31, *q* value 0.038) (Table S1), and only in zebrafish infected with the pneumolysin-deficient mutant strain, suggesting that the recognition of pneumolysin in early pneumococcal meningitis in zebrafish most likely depends on a different PRR pathway.

Necroptosis is a caspase-independent programmed form of necrosis that regulates inflammation and has been implicated to play an important role in infectious disease pathology (Kearney and Martin, 2017; Linkermann and Green, 2014; Vandenabeele et al., 2010). Interestingly, the core necroptotic machinery in mammals consists of RIPK3 and MLKL, the latter which is not present in zebrafish (Dondelinger et al., 2016). Nevertheless, treatment with necrostatin-5 or GSK’872 resulted in a higher mortality rate and higher bacterial load in infected zebrafish larvae (Fig. 2). This suggests that zebrafish have an alternative cell death executor instead of MLKL, or that RIPK3 triggers apoptosis. Alternatively, RIPK1-RIPK3 activation may restrict bacterial proliferation by a different pathway, for instance by triggering inflammation independent of MLKL (Orozco et al., 2019). Recent studies show that diverse bacterial pore-forming toxins, including pneumolysin, can also induce necroptosis in macrophages and lung epithelial cells (Gilley et al., 2016; González-Juarbe et al., 2015; Gonzalez-Juarbe et al., 2018; Kitur et al., 2015). Notably, we found that the necroptosis pathway was one of the most enriched pathways in zebrafish larvae injected with the *S. pneumoniae* PLY+ strain, indicating that pneumolysin plays a role in activating necroptosis in pneumococcal meningitis. Chemical inhibition of necroptosis resulted in higher mortality and bacterial burden in infected zebrafish larvae, suggesting a protective role for necroptosis. The role of necroptosis in infection is a topic of debate. The study of Kitur et al. showed that necroptosis limits pathological inflammation and enhances survival in a murine *S. aureus* skin and sepsis model (Kitur et al., 2016), whereas other studies have shown that pore-forming toxin-induced necroptosis exacerbates pulmonary injury and is detrimental to the host during bacterial pneumonia (González-Juarbe et al., 2015; Gonzalez-Juarbe et al., 2018; Kitur et al., 2015). Elucidating the role of necroptosis may provide new avenues for treatment of infectious diseases.

Finally, we provide the first visualization of pneumococcal competence development *in vivo* at the single cell level. This shows that, in contrast to the *in vitro* situation, where all cells within a clonal population synchronously become competent (Domenech et al., 2018; Martin et al., 2010; Moreno-Gámez et al., 2017; Slager et al., 2014), during meningitis in zebrafish, not all pneumococcal cells develop competence at the same time (Fig. 5A). It also challenges the idea that competence is constitutively expressed by individual bacteria in the host (Lin et al., 2020), but rather that there are pockets of pneumococci inducing competence during infection (Fig. 5). This is in line with evidence that cell-to-cell communication and cell-to-cell heterogeneity are of major importance in the pathogenesis of infectious diseases (Camilli and Bassler, 2006; Rutherford and Bassler, 2012). Competence in *S. pneumoniae* is regulated by a quorum sensing system, and is triggered by chemical signals and environmental factors such as antibiotic-induced replication stress, environmental cues, cell density and cell history (Moreno-Gámez et al., 2017; Slager et al., 2014). This system operates via the secreted peptide competence stimulating factor (CSP). Inactivation of the *comCDE* operon, which encodes the CSP precursor, histidine kinase and a response regulator results in complete abolishment of competence development (Slager et al., 2014). While competence allows for DNA uptake, most of the genes that are part of the competence regulon are not involved in transformation and likely function as a general stress response (Claverys et al., 2006; Engelmoer and Rozen, 2011). We observed that deletion of *comCDE* attenuated virulence in pneumococcal meningitis, which is in line with recent work that also showed a key role for competence in a murine meningitis model (Figure 5C-E; (Schmidt et al., 2019)). Interestingly, a recent study showed that to successfully infect the host, *S. pneumoniae* heterogeneously expresses pneumolysin to overcome host defense strategies and cross the blood-brain barrier by rising different bacterial subpopulations that can either attack or evade autophagosomes (Surve et al., 2018). Moreover, both competence as well as heterogeneity have been reported to play a role in streptococcal biofilm formation, an important virulence factor for nasopharyngeal colonization, pneumonia and otitis media (Cvitkovitch et al., 2003; Suntharalingam and Cvitkovitch, 2005; Vidal et al., 2013). This suggests that cell-to-cell communication and cell-to-cell heterogeneity in pneumococci is indeed involved in the pathogenesis of pneumococcal infection. It remains to be elucidated how the heterogeneous competence expression is involved, whether directly or indirectly, in pneumolysin-related virulence.

In summary, our work provides a detailed view of host-pathogen transcripts during early pneumococcal meningitis. The transcriptome data can serve as a valuable resource for future studies, with emphasis on genes and pathways we did not elaborate on in this study. Moreover, we made the dataset available online on https://veeninglab.com/dual-danio and encourage other researchers to use the dataset to further validate research findings and develop new hypotheses. Understanding the role and mechanism of pneumolysin in host-pathogen interactions will provide clues for future treatment strategies to combat not only pneumococcal meningitis but also other forms of invasive pneumococcal disease.

## DATA AVAILABILITY

The transcriptomics datasets are available in GEO repository, accession number GSE123988.

## ACKNOWLEDGMENTS

We thank Vladimir Benes (GeneCore, EMBL, Heidelberg) for his continuing support in library preparation and sequencing. We would like to acknowledge the Center for Information Technology of the University of Groningen for their support and for providing access to the Peregrine high-performance computing cluster. We thank Petr Broz (Biochemistry, University of Lausanne), Coen Kuijl (Medical Microbiology and Infection Prevention, Amsterdam UMC) and Astrid van der Sar (Medical Microbiology and Infection Prevention, Amsterdam UMC) for valuable discussions. We thank Annemarie Meijer (Institute of Biology, Leiden University) for providing the *Tg(cxcl18b:eGFP)* transgenic zebrafish line and Theo Verboom (Medical Microbiology and Infection Prevention, Amsterdam UMC) and Jeroen Kole (Confocal.nl) for technical support. We thank Doran Pauka for building the dual-danio website. Work in the Veening lab is supported by the Swiss National Science Foundation (SNSF) (project grants 310030_192517 and 310030_200792), a JPIAMR grant (40AR40_185533) from SNSF, NCCR ‘AntiResist’ from SNSF (51NF40_180541) and ERC consolidator grant 771534-PneumoCaTChER. D. van de Beek is supported by a ZonMw Vici grant (Vici 91819627).

## AUTHOR CONTRIBUTIONS

Conceptualization, K.K.J, R.A., W.B., and J-W.V.; bacterial strain creation, K.K.J, R.A., J.K., and A.D; dual RNA-seq sample preparation, K.K.J. and R.A., dual RNA-seq data analysis, K.K.J. and R.A. with input from W.B. and J-W.V.; experimental design, K.K.J., R.A., D.v.d.B., C.M.J.E.V-G., W.B., J-W.V.; manuscript writing, K.K.J, R.A., D.v.d.B., C.M.J.E.V-G., W.B., and J-W.V. with input from all authors.

## MATERIALS AND METHODS

### Bacterial strains and growth conditions

All pneumococcal strains used in this study are derivatives of the clinical isolate *S. pneumoniae* D39 from the Veening lab (D39V), unless specified otherwise and are listed in Table S4 (Avery et al., 1944; Slager et al., 2018). Oligonucleotides are listed in Table S5. *S. pneumoniae* was grown at 37°C on Columbia agar blood plates supplemented with 5% sheep blood (bioMérieux; 43049) or in C+Y medium (Martin et al., 1995). For transformation, pneumococcal cells were grown in C+Y medium at 37°C to an OD595 of approximately 0.100. Subsequently, 100 ng/ml of synthetic CSP-1 was added and the cells were incubated for 10 min at 37°C. DNA was added to the activated cells and incubated for 20 min at 30°C. Cells were then diluted 10 times in fresh C+Y medium and incubated for 1.5 h at 37°C. Transformants were selected by plating on Columbia agar blood plates supplemented with 2% (v/v) defibrinated sheep blood with the appropriate antibiotics. Antibiotic concentrations for selection used for *S. pneumoniae* were: erythromycin (ery) 0.25 μg/mL, chloramphenicol (cam) 4.5 μg/mL, kanamycin (kan) 250 μg/mL, spectinomycin (spec) 100 μg/mL, and trimethoprim (trmp) 20 μg/mL.

### Construction of pneumolysin deficient *S. pneumoniae* D39V PLY-mutant strain and its complemented *S. pneumoniae* D39V PLY+ strain

To construct the pneumolysin-deficient mutant strain PLY-, we transferred the pPEPY plasmid, which integrates at the chromosomal integration locus (CIL) in a non-coding region of the pneumococcal genome, to a previously published pneumolysin deficient mutant strain (Hassane et al., 2017; Keller et al., 2019), resulting in strain RA49. The isogenic complemented strain PLY-was constructed by ectopic integration of the PEPY plasmid with pneumolysin under native promoter and a kanamycin marker, resulting in strain LK01. This construct was made by amplifying the pneumolysin promoter from genomic *S. pneumoniae* D39V DNA with primer set LK283 and LK284. Pneumolysin with native terminator was amplified from genomic D39 DNA using primer set LK285 and LK286. Terminal primers LK283 and LK286 were used to join the two fragments together and contain restriction sites BamHI and XbaI respectively.

### Construction of fluorescently labelled S. *pneumoniae* D39V Δ*comCDE* mutant strain and its complemented strain

To construct the *comCDE* deficient mutant strain, erythromycin resistance marker (*ery*^*r*^) was amplified with using primers OVL2549 and OVL2771 from genomic DNA of the *hexA::ery*^*r*^ strain (Veening lab collection). The upstream region of *comCDE* was amplified using primers OVL506 and OVL2548, the downstream region with primers OVL2773 and OVL1667 using genomic DNA of *S. pneumoniae* D39V as template. The resulting three fragments were fused by Golden Gate cloning with BsmBI, and transformed into the complemented *S. pneumoniae* D39V PLY+ strain. Transformants were selected on Columbia blood agar containing 0.25 μg/mL of erythromycin, and correct colonies were verified by PCR and sequencing. To fluorescently label *S. pneumoniae* strains that have already chloramphenicol resistance marker (cam^r^), we replaced cam^r^ with trimethoprim resistance marker (trmp^r^) on *hlpA-sfGFP/RFP_cam*^*r*^ constructs. The *trmp*^*r*^ gene was amplified using primers OVL2549 and OVL2772 from genomic DNA of the *hexA::trmp*^*r*^ strain (Veening lab collection). Using genomic DNA of JWV500 (*hlpA::hlpA-sfGFP_cam*^*r*^) (Kjos et al., 2015) or MK119 (*hlpA::hlpA_hlpA-mmKate2_cam*^*r*^) (Kjos and Veening, 2014), the upstream fragment containing *hlpA-sfGFP or hlpA_hlpA-mmKate2* was amplified by PCR with primers OVL43 and OVL2769, and the downstream fragment of *cam*^*r*^ gene containing transcriptional terminator was amplified using primers OVL2770 and OVL46. The resulting three fragments were fused by Golden Gate cloning with *BsmB*I, and transformed into the respective *S. pneumoniae* strains, resulting in strains VL2725 and VL2728. Transformants were selected on Columbia blood agar containing 20 μg/mL of trimethoprim, and correct colonies were verified by PCR and sequencing.

### Construction of SsbB-GFP, HlpA-mCherry reporter strain

To monitor competence development of pneumococci *in vivo*, we transformed the amplified PCR product of the hlpA_hlpA-mCherry fragment and a downstream chloramphenicol cassette using chromosomal DNA of strain MK218 as a template into a strain containing a translational fusion of GFP to the competence-induced SsbB protein (Aprianto et al., 2016; Beilharz et al., 2015). This strain, KJ43, expresses constitutively mCherry and expresses SsbB-GFP upon competence induction. Transformants were selected on Columbia blood agar containing 4.5 μg/mL of chloramphenicol.

### Construction of Δ*cbpD* and isogenic Δ*comCDE*Δ*cbpD* mutant strains

To construct the Δ*cbpD* mutant strain, the gene *cbpD*, encoding the fratricin called choline-binding protein D, was replaced by a spectinomycin resistance (spec^r^) marker. The upstream region was amplified using primers ADP1/45 and ADP1/46+SphI, the downstream region with primers ADP1/47+HindIII and ADP1/48, and the spectinomycin resistance marker with sPG11+SphI and sPG12+HindIII. All three fragments were digested with the proper restriction enzymes (SphI and/or HindIII) and ligated together. The Δ*cbpD*::spec fragment containing the spectinomycin resistance marker flanked by the sequence up- and downstream of *cbpD* was transformed into *S. pneumoniae* D39V resulting in strain VL561. Transformants were selected on Columbia blood agar containing 200 μg/mL spectinomycin. Correct deletion was verified by PCR and sequencing. To construct the isogenic Δ*comCDE*Δ*cbpD* mutant strain, the Δ*comCDE*::cat fragment from strain ADP107 (Moreno-Gámez et al., 2017)was transformed into VL561 resulting in strain ADP351. All transformants were selected on Columbia blood agar containing 3.5 μg/mL of chloramphenicol, and correct colonies were verified by PCR and sequencing.

### Zebrafish husbandry and maintenance

Transparent adult *casper* mutant zebrafish (D’Agati et al., 2017; White et al., 2008)and *Tg(cxcl18b:eGFP)* (Torraca et al., 2017) wild-type zebrafish expressing green fluorescence upon induction of the inflammatory marker chemokine cxcl18b were maintained at 26°C in aerated 5-L tanks with a 10/14 h dark/light cycle. Zebrafish embryos were raised at 28°C in a temperature-controlled incubator in E3 medium (5.0 mM NaCl, 0.17 mM KCl, 0.33 mM CaCl_2_·2H_2_O, 0.33 mM MgCl_2_·6H_2_O) supplemented with 0.3 mg/L methylene blue. If necessary, *Tg(cxcl18b:eGFP)* wild-type zebrafish were additionally treated with 0.003 % (v/v) 1-phenyl 2-thiourea (PTU) to inhibit the formation of melanocytes (Karlsson et al., 2001). All procedures involving zebrafish embryos were according to local animal welfare regulations.

### Simultaneous total host-pathogen RNA isolation

*Casper* zebrafish larvae were injected with similar doses of the pneumolysin deficient *S. pneumoniae* D39V (PLY-) mutant strain or the isogenic complemented *S. pneumoniae* D39V (PLY+) strain at 2 dpf as described below. At 8 hours post injection (hpi), 100 individually infected larvae per group were anesthetized with 0.02% Tricaine, pooled into one biological replicate. In total 3 biological replicates per group (n=6) were isolated. To minimize transcriptional changes during sample handling, samples were snap-frozen in liquid nitrogen immediately after the study was completed. Further, we simultaneously harvested host and pathogen cells and isolated combined total RNA. To harvest the total RNA, we treated the infected larvae with a concentrated solution of ammonium sulfate to prevent any protein-dependent RNA degradation (Korfhage et al., 2002). Each milliliter of the ammonium sulfate solution (pH 5.2) contained 0.7 g (NH_4_)_2_SO_4_. The saturated solution also contained 20 Mm EDTA and 25 mM sodium citrate. Three parts of saturated solution of ammonium sulfate was added directly to one part of medium. The suspension was vigorously pipetted to ensure the complete mixing of the ammonium sulfate solution and the samples, and incubated further (room temperature, 5 min). The suspension was collected and centrifuged at full speed (20 min, 4°C, 10,000 × *g*). The supernatant was removed and the cell pellet was snap-frozen with liquid nitrogen.

In a 1.5 mL screw cap tube, a PCR tube full of sterile, RNase-free glass beads (100 μm, BioSpec, US) were added together with 50 μL10% SDS and 500 μL phenol-chloroform. The frozen pellet was resuspended in TE solution (10 mM Tris-HCl, 1 mM Na_2_DTA, pH 8.0, DEPC treated mQ). The cell suspension was added into the screw cap tube and bead-beaten three times, 45 s each. Tubes were immediately placed on ice and centrifuged at full speed at 4°C to separate organic and aqueous phases. The aqueous phase was pipetted out and back-extraction was performed on the organic phase to optimize RNA yield. Phenol was further depleted from the aqueous phase by a round of chloroform extraction. After vigorous vortexing, the mixture was again centrifuged at full speed, 4°C. The aqueous phase was pipetted into a new Eppendorf tube and nucleic acids were alcohol-precipitated: 50 μl NaOAc 3M and 1 mL of cold isopropanol were added and mixed thoroughly. The mixture was incubated for at least 30 min at −20°C before pelleting by centrifugation (full speed, 4°C). Supernatant was removed gently and the nucleic acid pellet was resuspended in ice-cold 75% ethanol before re-pelleting was performed (full speed, 4°C). Ethanol washing was performed once more. The pellet was air-dried before DNase treatment.

DNase treatment was performed according to the manufacturer’s protocols (RNase-free DNase I recombinant, Roche, US) for 1 hour, room temperature. To remove DNase and gDNA-derived nucleotides, phenol-chloroform extraction, chloroform extraction, isopropanol precipitation and ethanol washing were performed as previously mentioned. Total RNA was resuspended in 30 μl TE buffer. The quantity and quality of total RNA was estimated by Nanodrop and a 1% bleach gel (Aranda et al., 2012) was employed to interrogate the presence of genomic DNA and rRNA bands (23S, 2.9 kbp and 16S, 1.5 kbp).

### Library preparation and sequencing

Total RNA was isolated and the RNA quality checked using chip-based capillary electrophoresis. Samples were simultaneously depleted from eukaryotic and Gram positive ribosomal RNAs by dual rRNA-depletion as previously described (Aprianto et al., 2016). Stranded cDNA library preparation with the TruSeq® Stranded Total RNA Sample Preparation Kit (Illumina, US) according to the manufacturer**’**s protocol. Sequencing was performed for twelve samples in one lane of Illumina NextSeq 500, High Output Flowcell in 85 single end mode. Libraries were demultiplexed and analyzed further. Raw libraries are accessible at https://www.ncbi.nlm.nih.gov/geo/ with the accession number GSE123988.

### Data analysis

Quality of raw libraries was checked (Andrews, 2010) (FastQC v0.11.8, Babraham Bioinformatics, UK). In order to improve the quality of alignment, we trimmed the reads using the following criteria: (i) removal of adapter sequence, if any, based on TruSeq3-SE library, (ii) removal of low quality leading and trailing nucleotides, (iii) a five-nucleotide sliding window was created for surviving reads, in which the average quality score must be above 20 and (iv) minimum remaining length must be above 50 (Trimmomatic v0.38) (Bolger et al., 2014). The quality of trimmed reads was confirmed using FastQC (Andrews, 2010).

We opted to align the reads to a chimeric genome containing the concatenated circular genome of *S. pneumoniae* onto the zebrafish genome. Here, we created chimeric genomes by concatenating the circular genome of *S. pneumoniae* strain D39V (GCF_003003495) onto the genome of *Danio rerio* (ftp://ftp.ensembl.org/pub/release-94/fasta/danio_rerio/dna/, accessed 13 November 2018). The corresponding annotation file was downloaded from the same ftp folders. Alignment was performed by RNA-STAR (v2.6.0a) (Dobin et al., 2013) with the following options: (i) alignIntronMax 1 and (ii) sjdbOverhang 80. The aligned reads were the summarized (featureCount v1.6.3) according to the chimeric annotation file in stranded, multimapping (-M), fractionized (--fraction) and overlapping (-O) modes^7^. The single-pass alignment was selected onto chimeric genome was selected to minimize false discovery rate.

We then analyzed host and pathogen libraries separately in R (R v3.5.2). Because rRNA depletion was successful (relative pneumococcal rRNA count ∼ 2.18%, relative zebrafish rRNA count ∼0.4%), we did not exclude reads which aligned to genes encoding ribosomal RNA in subsequent analysis. Differential gene analysis was performed by DESeq2 v1.22.1 (Love et al., 2014)and genome-wide fold change was calculated between the transcriptional response of the two bacterial strains, PLY+ and PLY-. Value of fold change was set to zero if the corresponding *q* value is reported to be NA. Enrichment tests for functional analysis were performed by the built-in function, *fisher*.*test()*.

Soft clustering was performed on the normalized centered gene expression values with a fuzzifier value of 1.5 to obtain a better view of the dynamics of gene expression during infection (Kumar and Futschik, 2007). Functional enrichment analysis was performed using WEB-based GEne Set AnaLysis Toolkit (WebGestalt) (Liao et al., 2019). For Gene Ontology (GO) enrichment analysis at least 3 genes were required and *p* value < 0.05. Gene Set Enrichment Analysis was analyzed by Webgestalt 2019

### qRT-PCR confirmation of host genes

To obtain total RNA, we first anesthetized zebrafish larvae (n=50 per group) in 0.02% Tricaine in egg water at 8 hpi and collected them in a 2 mL Eppendorf tube. Then, excess egg water was removed and 350 μl lysis buffer added (NucleoSpin, MACHEREY-NAGEL). The larvae were homogenized by repeatedly drawing through a 26-gauge needle into a syringe and expelled. Isolation of total RNA was performed using NucleoSpin RNA isolation kit (NucleoSpin, MACHEREY-NAGEL). cDNA was synthesized from total RNA by SuperScript® IV Reverse Transcriptase (Life Technologies, NL). Primers were designed across exons (for host genes) while retaining its species specificity as confirmed by *in silico* PCR on the opposite species (i.e., host primers to *S. pneumoniae* genome, *vice versa*) (Table S5). Amplification efficiency for each primer was calculated based on primer ability to double amount of product per cycle. The qPCR mix contained forward and reverse primers, PowerTrack™ SYBR Green Master Mix (Applied Biosciences; A46012) and cDNA. The reactions were on triplicates and two different cDNA concentrations. Subsequently, PowerTrack™ SYBR Green Master Mix and primers for the selected genes (Table S5) were added and RT-qPCR was performed on a StepOnePlus™ Real-Time PCR System (Applied Biosystems). As reference genes we used the *Mobk13 (mob4)* for zebrafish genes and *gyrA* for pneumococcal genes (Aprianto et al., 2016; Hu et al., 2016). The RT-qPCR fold change was calculated by the ΔΔCt method (Livak and Schmittgen, 2001).

### Fluorescence imaging

Screening and imaging of *Tg(cxcl18b:eGFP)* zebrafish larvae were performed with a Zeiss Axio Zoom V16 stereo microscope with a Hamamatsu ORCA flash 4.0 camera attached. Zebrafish larvae infected with the *S. pneumoniae* D39V *ssbB*-reporter strain were collected at 8 hours post injection and fixated overnight in 4 % paraformaldehyde in PBS, washed with PBS, prior to confocal imaging. Confocal images were generated with a Rescan Confocal Microscope 2 (RCM2, Confocal.nl) with a Hamamatsu ORCA flash 4.0 v3. For optimal imaging, embryos were embedded in 1.5 % low-melting-point agarose dissolved in PBS in an open uncoated 8-well microscopy μ-Slide (http://ibidi.com). ImageJ software was used to process the confocal images, specifically for brightness/contrast enhancements as well as for creating merged images.

### Zebrafish survival experiments

Pneumolysin deficient *S. pneumoniae* D39V PLY-mutant strain or complemented *S. pneumoniae* D39V PLY+ strain were grown in C + Y medium until mid-log phase (OD595nm ∼0.2), harvested by centrifugation (6000 RPM for 10 min.), washed with sterile PBS, harvested again by centrifugation, and finally resuspended in 0.125 % (w/v) phenol red solution (Sigma-Aldrich; P0290) to aid visualization of the injection process. Prior to injection, 2 days post fertilization (dpf) embryos were mechanically dechorionated if necessary and anaesthetized in 0.02 % (w/v) buffered 3-aminobenzoic acid methyl ester (pH 7.0) (Tricaine; Sigma-Aldrich, A5040). The bacterial suspension was then injected into the hindbrain ventricle of *casper* zebrafish larvae at 2 dpf. After injection, the larvae were kept in 6-well plates containing egg water (60 μg/mL sea salts (Sigma-Aldrich; S9883) in demi water) at 28 °C and the mortality rate monitored at fixed time-points as described previously (Jim et al., 2016). To assess the effect of necroptosis pathway inhibition on survival, infected zebrafish larvae were treated orally with 100 μM necrostatin-5 (Sigma-Aldrich; N0164), a RIPK1 inhibitor, GSK’872 (MedChemExpress; HY-101872), a RIPK3 inhibitor or DMSO (vehicle control) by adding the compounds to egg water. All experiments were performed in triplicate. Survival graphs were generated with GraphPad Prism 8.0 and analyzed with the log rank (Mantel-Cox) test. Results were considered significant at *p* values <0.05.

### Bacterial load and competitive index

To determine the bacterial load at different time-points, infected zebrafish larvae were anesthetized in 0.02 % Tricaine in egg water, transferred to a 1.5 mL screwcap tube (1 larva per tube) filled with 1.0 mm glass beads (Sigma-Aldrich, Z250473) to ∼25% capacity of the tubes’ volume, placed in a microvial rack, and violently shaken (3 times 10 sec, 10 sec interval) in a bead beater (BioSpec Products; Mini Beadbeater) to disrupt the cells and tissues. Subsequently, serial dilution plating was performed on Columbia Blood Agar (BD Biosciences, 211124) plates supplemented with 5% defibrinated sheep blood (bioTRADING, BTSG100) (COS plates), 10 mg/L colistin sulphate and 5 mg/L oxolinic acid (COBA, Oxoid), to inhibit growth of commensal bacteria in zebrafish. The plates were incubated O/N at 37°C and quantified the next day. To determine the competitive indexes, zebrafish larvae were co-injected at 2 dpf with two pneumococcal strains of interest mixed in a 1:1 ratio into the hindbrain ventricle. The number of colony-forming units (CFU) per injection was determined by quantitative plating of the injection volume on COS plates containing appropriate antibiotics. At 24 hpi, anesthetized larvae were homogenized as described above and serial dilution plating was performed on COS plates containing COBA supplement and appropriate antibiotics, to quantify the number of both pneumococcal strains in each larva. The competitive indexes were then calculated by dividing the ratio of the test strain divided by the reference strain. All experiments were performed in duplicate. Scatter plot and competitive index plots were generated with GraphPad Prism 8.0 and analyzed with unpaired t-test or one sample t-test respectively. Results were considered significant at *p* values <0.05.

## SUPPLEMENTAL FIGURES

**Figure S1.**
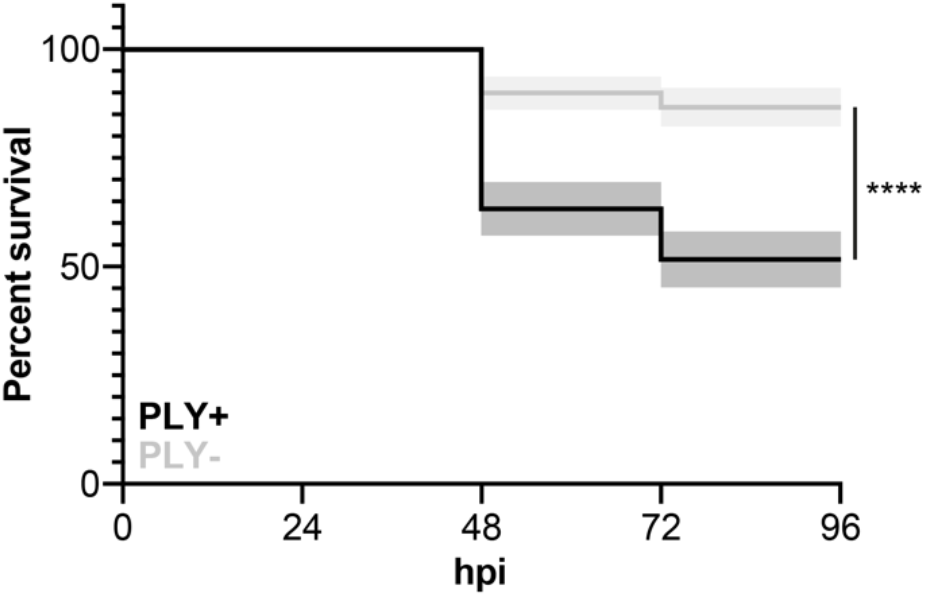
Pneumolysin-deficient *S. pneumoniae* strain is attenuated in the zebrafish pneumococcal meningitis model. Survival curves of 2 dpf zebrafish larvae injected with *S. pneumoniae* D39V PLY+ or *S. pneumoniae* D39V PLY-in the hindbrain ventricle. Larvae were infected with 400 CFU. The data represent the mean ± SEM of three individual experiments with 20 embryos in each group; *p* value < 0.0001; determined by log-rank test.

**Figure S2.**
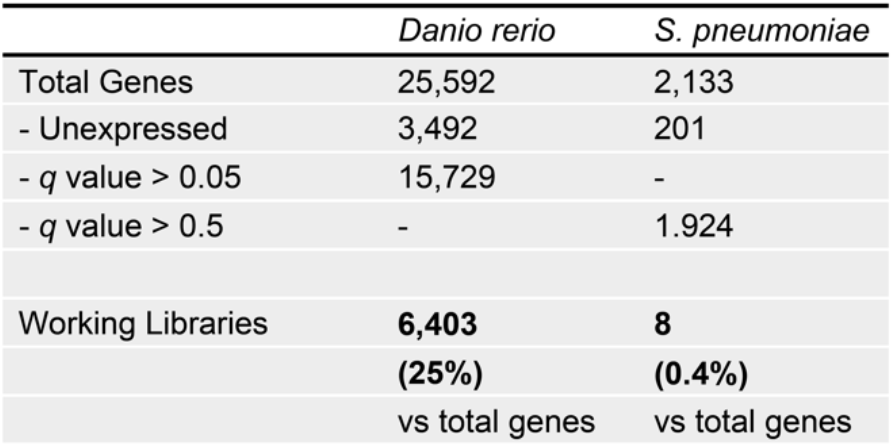
Zebrafish and pneumococcal working libraries. Two gene fractions were removed to simplify downstream analysis. After removal of unexpressed genes and non-significant genes the zebrafish working library contained 6403 genes (25% of *D. rerio* genes) whereas the pneumococcal working library contained 8 genes (0.4% of *S. pneumoniae* genes).

**Figure S3.**
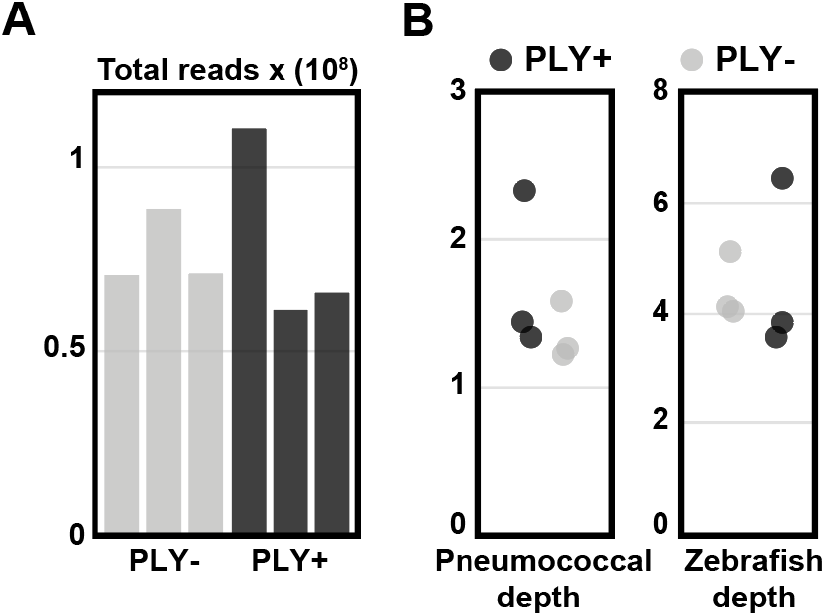
Total reads and depth. **(A)** On average, each library has 77 million reads (467 million reads in total); equivalent to 6.2 billion nucleotides. **(B)** Average depth of pneumococcal reads is 1.54 and zebrafish depth is 4.54.

**Figure S4.**
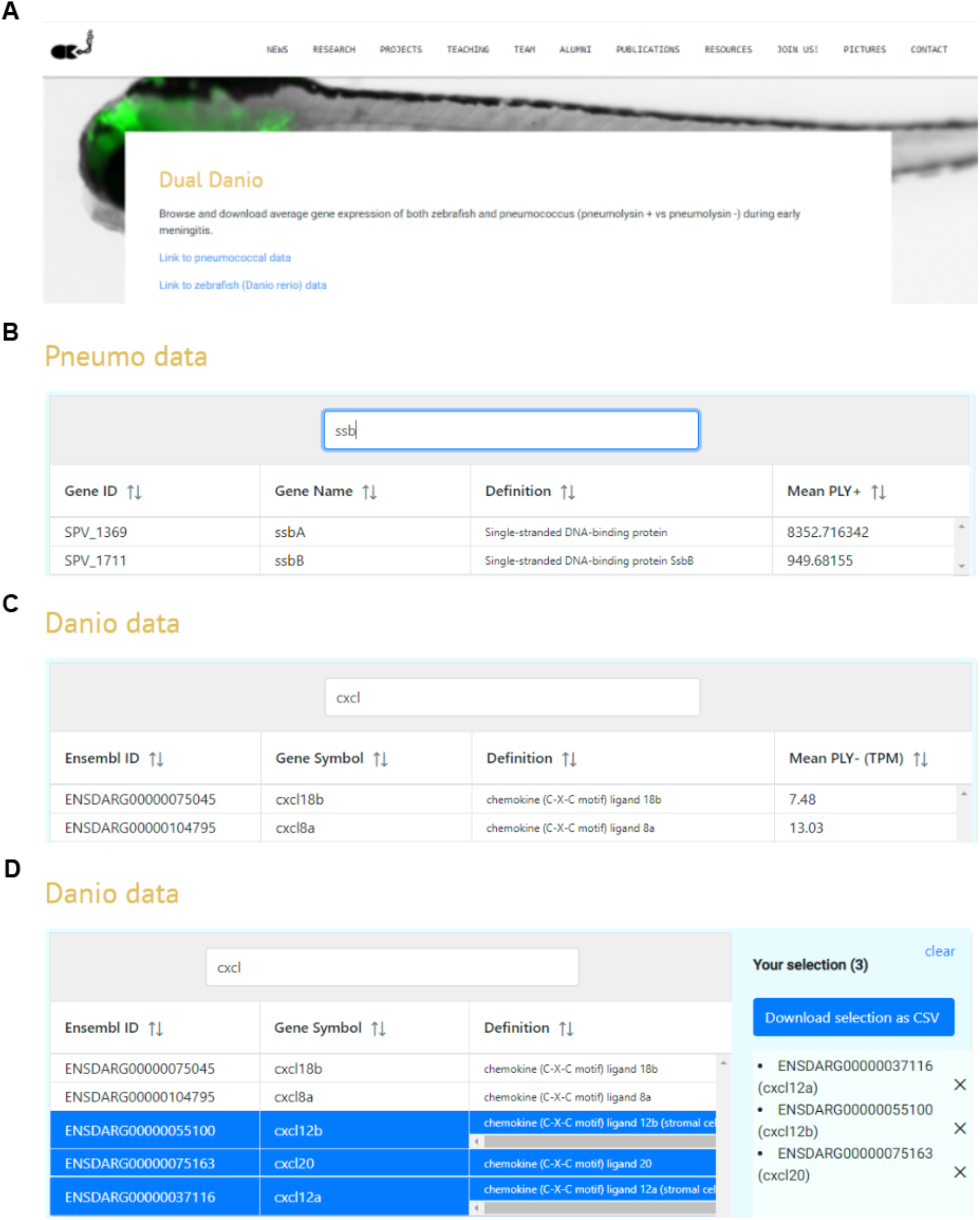
Easy access of the complete dual RNA-seq database. **(A)** Screenshot of the web-based platform with search result example summary for *D. rerio* **(B)** or *S. pneumoniae* **(C)**. The RNA-seq data of a single gene or multiple genes can be easily exported **(D)**.

## SUPPLEMENTARY TABLES

**Table S1**. *Danio rerio* differential gene expression

**Table S2**. Functional enrichment analysis of differentially expressed *Danio rerio* genes

**Table S3**. *Streptococcus pneumoniae* D39V differential gene expression

**Table S4**. Strains

**Table S5**. Primers

## SUPPLEMENTARY MOVIES

**Movie S1**. Cxcl18b expression over time in infected versus control zebrafish larva

